# Contrasting dynamics of two incursions of low pathogenicity avian influenza virus into Australia

**DOI:** 10.1101/2024.04.23.590662

**Authors:** Michelle Wille, Ivano Broz, Tanya Cherrington, Allison Crawley, Blaine Farrugia, Mark Ford, Melinda Frost, Joanne Grimsey, Peter D. Kirkland, Shaylie Latimore, Stacey E. Lynch, Sue Martin, Cornelius Matereke, Peter T. Mee, Matthew J. Neave, Mark O’Dea, Andrew J. Read, Kim O’Riley, Vittoria Stevens, Sivapiragasam Thayaparan, Sara Zufan, Silvia Ban de Gouvea Pedroso, Victoria Grillo, Andrew C. Breed, Ian G. Barr, Edward C. Holmes, Marcel Klaassen, Frank Y. K. Wong

## Abstract

The current panzootic of high pathogenicity avian influenza virus H5N1 demonstrates how viral incursions can have major ramifications for wildlife and domestic animals. Herein, we describe the recent incursion into Australia of two low pathogenicity avian influenza virus subtypes, H4 and H10, that exhibited contrasting evolutionary dynamics. Viruses detected from national surveillance and disease investigations between 2020-2022 revealed 27 genomes, 24 of which have at least one segment more closely related to Eurasian or North American avian influenza lineages than those already circulating in Australia. Phylogenetic analysis revealed that H4 viruses circulating in shorebirds represent a recent incursion from Asia that is distinct from those circulating concurrently in Australian waterfowl. Analysis of the internal segments further demonstrates exclusive, persistent circulation in shorebirds. This contrasts with H10, where a novel lineage has emerged in wild waterfowl, poultry and captive birds across Australia, and has likely replaced previously circulating H10 lineages through competitive exclusion. Elucidating different dynamics for avian influenza incursions supports effective disease risk identification and communication that better informs disease preparedness and response.

## Introduction

High pathogenicity avian influenza virus (HPAIV) H5N1 is causing a major disease burden on the poultry industry and in wild birds and marine mammals (Wille et al. 2022). The current panzootic encompasses Europe, Asia, Africa, North America, South America and Antarctica; with Oceania the only continent still free from HPAI H5N1 (Wille et al. 2024). Despite the negative consequences for wild bird populations, migratory avian hosts facilitate long-distance virus dispersal of HPAIV H5N1, a role that has increased over the years with the emergence of HPAIV H5Nx clade 2.3.4.4 in 2014 and 2.3.4.4b in 2021 (Global Consortium for H5N8 and Related Influenza Viruses 2016, Wille et al. 2022, European Food Safety Authority et al. 2023, Klaassen et al. 2023). Improved understanding of avian-borne viral movement and incursions has therefore become increasingly important to improved disease preparedness and in mounting appropriate responses.

Beyond HPAIV H5N1, wild birds are the natural reservoirs for low pathogenicity avian influenza viruses (LPAIV). Wild birds, particularly members of the orders *Anserifomes* (waterfowl, including ducks, geese, swans) and *Charadriiformes* (shorebirds and gulls) are principal reservoirs for LPAIV, with 16 HA (haemagglutinin) and 9 NA (neuraminidase) subtypes identified to date (Olsen et al. 2006). These host orders play differing roles in LPAIV ecology, with contrasting patterns of prevalence (Wille et al. 2023), seasonal ecology (Maxted et al. 2016), and roles in long-distance dispersal between continents (Rimondi et al. 2018, Wille et al. 2022), which is clearly apparent in the Australian context (e.g. (Hansbro et al. 2010, Hoque et al. 2015, Hoye et al. 2021, Wille et al. 2022). In addition, there is relatively little transfer of viral lineages between these host groups, although it does occur (Hicks et al. 2022). In contrast to waterfowl from other continents, Australian wild waterfowl are endemic to the Australo-Papuan region and do not connect Australia with Eurasia and North America through migration (McCallum et al. 2008). Rather, long-distance migratory shorebirds annually migrate between Australia and breeding areas in Eastern Siberia and Alaska, with key stop-over sites across Asia (Tracey et al. 2004, McCallum et al. 2008). As such, viral introductions to Australia are likely facilitated by long-distance migratory shorebirds rather than waterfowl, with intercontinental reassortment a feature of some viral genomes from shorebirds sampled in Australia (Hurt et al. 2006, Hoye et al. 2021, Wille et al. 2022). Following these infrequent introductions, the persistent circulation of LPAIV lineages has been observed within the continent, presumably by nomadic waterfowl. As such, Australia can generally be regarded as a sink for virus diversity (Wille et al. 2022).

In this study, we analysed recently-detected LPAIV genomes associated with two recent viral introductions to better understand incursions to, and dispersal within Australia. Through time-scaled phylogenetic analysis we estimated the dates and probable origins of these viral introductions and illustrated the contrasting patterns of viral maintenance in wild birds following these events.

## Methods

### Ethics Statement

Capture, banding and sampling of shorebirds and ducks was conducted under Australian Bird and Bat Banding Scheme authorities 2915 and 8001. Animal ethics permits were provided by Deakin University Animal Ethics permit number B39-2019 and Philip Island Nature Parks Animal Ethics permit number SPFL20082. Faecal environmental surveillance samples collected by Department of Primary Industries, Parks, Water and Environment, Tasmania and Primary Industries and Regions, South Australia, Department of Primary Industries and Regional Development, Western Australia; NSW Department of Primary Industries did not require permits. Samples collected for disease investigations, similarly, did not require permits.

### Sample collection and screening

Sample collection through the National Avian Influenza in Wild Birds (NAIWB) program was undertaken as per Grillo *et al*. 2015 between 2020 and 2022, including samples from wild caught birds, hunted birds, and fresh faeces from the environment. In addition, we included passive surveillance samples from affected domestic or captive birds as part of diagnostic investigations (Table S1). Viral screening and sequencing was undertaken as per Wille *et al*. 2022. Briefly, RNA was extracted from swab samples in virus transport media and screened by real-time RT-PCR using primers and probes targeted against the influenza A virus matrix gene. Either original samples (with Ct <30) or viral isolates or both were sequenced on an Illumina MiSeq with up to 24 samples pooled per sequencing run by use of dual-index library preparation and the Nextera XT DNA Library Preparation kit and 300-cycle MiSeq Reagent v2 kit (Illumina). Sequence reads were trimmed for quality and mapped to respective reference sequence for each influenza A virus gene segment using Geneious Prime software (www.geneious.com) (Biomatters, Auckland, NZ) (Wille et al. 2022).

Sequences generated in this study have been deposited in GenBank (Accession numbers available in Table S1)

### Phylogenetic analysis

All full length H4, H10, N1, N2, N6, N8 sequences in the database were downloaded from the Bacterial and Viral Bioinformatics Resource Centre Influenza portal (BV-BRC; www.bv-brc.org) for the construction of global maximum likelihood trees. H10 genome sequences reported in (Vijaykrishna et al. 2013) were also downloaded from GISAID (https://gisaid.org/). Sequences were aligned using MUSCLE v3.8.425 (Edgar 2022) integrated within Geneious Prime. Maximum likelihood trees incorporating the best-fit model of nucleotide substitution estimated using Smart Model Selection (Lefort et al. 2017), and aBayes (Anisimova et al. 2011) node support were estimated in PhyML v3.0 (Guindon et al. 2010).

Prior to the estimation of time-scaled phylogenies we evaluated the extent of molecular clock-like structure in the data by performing linear regressions of root-to-tip distances against year of sampling using maximum likelihood trees using TempEst (Rambaut et al. 2016). As evidence for a molecular clock was obtained, time scaled phylogenetic trees were estimated using BEAST v1.10.4 (Drummond et al. 2012), under the relaxed uncorrelated lognormal relaxed clock (Li et al. 2012) and SRD06 codon structured nucleotide substitution model (Shapiro et al. 2006), with the exception of M and NS segments for which we used the HKY+G model due to overlapping reading frames. The Bayesian skyline coalescent tree prior was used as this likely reflects the complex epidemiology dynamics of avian influenza viruses through time (Drummond et al. 2005). 100 million generations were performed, and convergence was assessed using Tracer v1.8 (http://tree.bio.ed.ac.uk/software/tracer/). Maximum credibility lineage trees were generated using TreeAnnotator following the removal of 10% burnin, and trees were visualised using Fig Tree v1.4 (http://tree.bio.ed.ac.uk/software/figtree/).

## Results

### Virus detection

Both H4 and H10 viruses were recovered through the NAIWB targeted surveillance program and other *ad hoc* sampling between December 2020 and September 2022. Briefly, H4 viruses were recovered from hunter-shot ducks in Tasmania (n=1), and South Australia (n=2), faecal environmental samples in South Australia (n=1) a hunter-shot duck (*Anas gracilis*, n=1) in New South Wales and live-captured shorebirds (*Calidris* spp, n=5) in Victoria. H10 viruses were recovered from faecal environmental samples collected in South Australia (n=2) and Western Australia (n=1), and from live-captured ducks (*Anas* sp) in Victoria (n=10) (Table S1).

Additional H10 viruses were identified through passive surveillance events in domestic or captive birds by respective state government animal health laboratories in conjunction with the national reference laboratory at the CSIRO Australian Centre for Disease Preparedness. These occurred between June-August 2021 and included chickens (*Gallus domesticus*) from New South Wales (NSW), farmed emu (*Dromaius novaehollandiae*) in Victoria, and captive Tawny Frogmouth (*Podargus strigoides*) in Western Australia (WA) (Table S1). The detection of an H10N7 virus from a chicken in New South Wales involved a backyard chicken farm approximately 20 km from Canberra, Australian Capital Territory. The flock of 234 birds experienced a morbidity of 12% and mortality of 6%. Movement restrictions were placed on birds from this farm with subsequent testing no longer able to detect avian influenza virus. In Victoria, an H10N3 virus was detected in farmed emu chicks aged three to six weeks with respiratory signs and low-level mortalities. Finally, in Western Australia, eight aviary-kept Tawny Frogmouths died suddenly at a Perth wildlife park, one of which was submitted for diagnostic laboratory investigation resulting in the detection of an H10N7 virus.

### Incursion and export events with limited host range of H4 viruses from shorebirds

H4 viruses recovered from waterfowl and shorebirds had different evolutionary histories, with viruses recovered from waterfowl falling into an H4 lineage which has persisted in Australia for four decades, compared to viruses from shorebirds which fell into a different lineage, comprising a recent incursion event (Figure 1, Figure S1).

**Figure 1.**
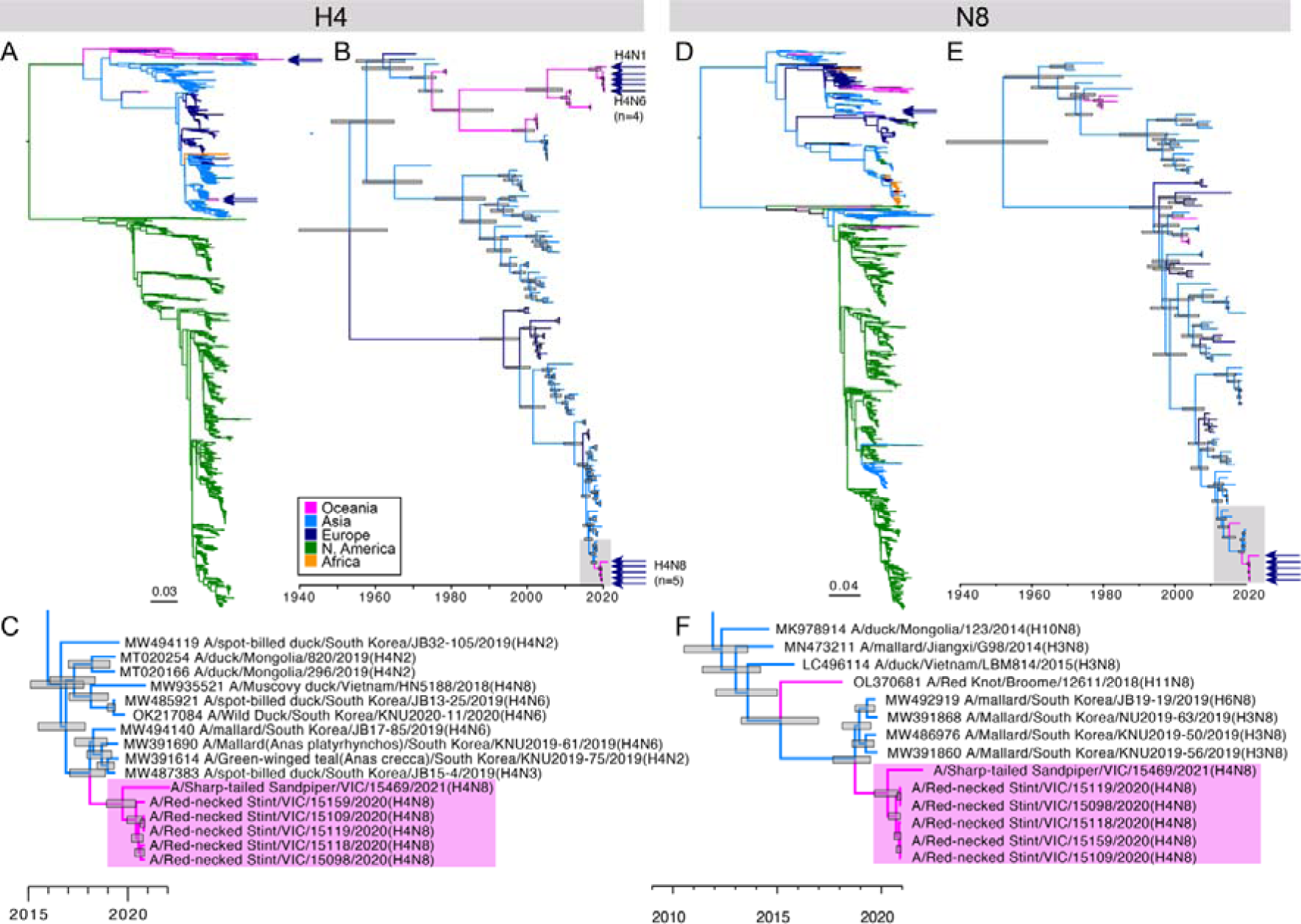
Phylogenetic trees of (A,B,C) H4 sequences regardless of NA, and (D,E,F) N8 sequences. Phylogenetic trees for the waterfowl clade of H4, and H4 associated NA trees are provided in Figure S1 and S2, respectively. (A, D) Maximum likelihood trees comprising all sequences collated for this study. Trees were rooted geographically, and the scale bar corresponds to number substitutions per site. (B, E) Time structured phylogenetic trees. The trees comprise select sequences from relevant lineages (here Eurasian lineages containing Australian sequences). (C, F), Expansion of clade containing Australian sequences of interest, highlighted in a grey box in (B, E). In the case of the HA segment, only the clade containing H4N8 viruses have been highlighted in C. Sequences of interest are indicated by arrows in A, B, D, E and are highlighted in a box in C, F. Node bars correspond to the 95% highest posterior density (HPD) of node height. Branches are coloured by continent. ches are coloured by continent.

Viruses from waterfowl included H4N6 (n=4) and H4N1 (n=1), with six of the eight gene segments of these viruses falling into established clades circulating in Australia (Table 1, Figure 1, S2-S3). The HA sequences of all five viruses fell into the same clade, despite being recovered from different Australian states (Figure 1). The N6 sequences similarly all belonged to a single clade, and for both N1 and N6 the sequences were most similar to other viruses that have circulated in Australia since 2019 (Figure S2). The internal gene constellations were largely similar, with PB1, PA, M and NP segments of all five H4 viruses belonging to phylogenetic clades previously detected in Australia. However the two H4N6 viruses detected in South Australia had PB2 and NS segments most closely related to sequences from Asia, and these viruses are likely the result of a reassortment event with undetected, recently introduced viruses of Eurasian lineage (Table 1, Figure 1, S2-S3).

**Table 1.** Viruses sequenced in this study and lineage delineations for all genome segments.

| Subtype | Designation | Sample date | PB2<sup>1</sup> | PB1<sup>1</sup> | PA<sup>1</sup> | HA<sup>1</sup> | NP<sup>1</sup> | NA<sup>1</sup> | M<sup>1</sup> | NS<sup>1</sup> |
| --- | --- | --- | --- | --- | --- | --- | --- | --- | --- | --- |
| H4N1 | A/Pacific Black Duck/Tasmania/22-1184-5/2022(H4N1) | 5/3/2022 | EUR - AUS (A) | EUR - AUS (A) | EUR - AUS (A) | EUR - AUS (H4) | NAm - AUS | EUR - AUS (N1) | EUR - AUS (A) | A-EUR - AUS (A) |
| H4N6 | A/Grey Teal/New South Wales/14340X/2020(H4N6) | 2/12/2020 | EUR - AUS (A) | EUR - AUS (A) | EUR - AUS (A) | EUR - AUS (H4) | NAm - AUS | EUR - AUS (N6) | EUR - AUS (A) | A-EUR - AUS (B) |
| H4N6 | A/Grey Teal/South Australia/22-64233564-20/2022(H4N6) | 21/5/2022 | EUR - novel | EUR - AUS (A) | EUR - AUS (A) | EUR - AUS (H4) | NAm - AUS | EUR - AUS (N6) | EUR - AUS (A) | A-EUR - novel |
| H4N6 | A/Grey Teal/South Australia/22-64233564-49/2022(H4N6) | 21/5/2022 | EUR - novel | EUR - AUS (A) | EUR - AUS (A) | EUR - AUS (H4) | NAm - AUS | EUR - AUS (N6) | EUR - AUS (A) | A-EUR - novel |
| H4N6 | A/wild waterbird/South Australia/22-68204541-55/2022(H4N6) | 16/8/2022 | na<sup>2</sup> | na<sup>2</sup> | na<sup>2</sup> | EUR - AUS (H4) | NAm - AUS | EUR - AUS (N6) | EUR - AUS (A) | B-EUR-AUS |
| H4N8 | A/Red-necked Stint/Victoria/15098/2020(H4N8) | 9/12/2020 | EUR - AUS (B) | EUR - AUS (B) | EUR - AUS (B) | EUR - novel (H4) | EUR - AUS | EUR - novel (N8) | EUR - AUS (B) | A-EUR - AUS (C) |
| H4N8 | A/Red-necked Stint/Victoria/15109/2020(H4N8) | 9/12/2020 | EUR - AUS (B) | EUR - AUS (B) | EUR - AUS (B) | EUR - novel (H4) | EUR - AUS | EUR - novel (N8) | EUR - AUS (B) | A-EUR - AUS (C) |
| H4N8 | A/Red-necked Stint/Victoria/15118(H4N8) | 9/12/2020 | EUR - AUS (B) | EUR - AUS (B) | EUR - AUS (B) | EUR - novel (H4) | EUR - AUS | EUR - novel (N8) | EUR - AUS (B) | A-EUR - AUS (C) |
| H4N8 | A/Red-necked Stint/Victoria/15119/2020(H4N8) | 9/12/2020 | EUR - AUS (B) | EUR - AUS (B) | EUR - AUS (B) | EUR - novel (H4) | EUR - AUS | EUR - novel (N8) | EUR - AUS (B) | A-EUR - AUS (C) |
| H4N8 | A/Red-necked Stint/Victoria/15159/2020(H4N8) | 9/12/2020 | EUR - AUS (B) | EUR - AUS (B) | EUR - AUS (B) | EUR - novel (H4) | EUR - AUS | EUR - novel (N8) | EUR - AUS (B) | A-EUR - AUS (C) |
| H4N8 | A/Sharp-tailed Sandpiper/VIC/15469/2021(H4N8) | 29/12/2021 | EUR - AUS (B) | EUR - AUS (B) | EUR - AUS (B) | EUR - novel (H4) | EUR - AUS | EUR - novel (N8) | EUR - AUS (B) | A-EUR - AUS (C) |
| H10N2 | A/Grey Teal/South Australia/22-64233564-37/2022(H10N2) | 21/5/2022 | EUR - AUS (A) | EUR - AUS (A) | EUR - AUS (A) | NAm - novel (H10) | NAm - AUS | EUR - novel (N2) | EUR - AUS (A) | B-EUR-AUS |
| H10N3 | A/emu/Victoria/21-03712/2021(H10N3) | 19/8/2021 | EUR - novel | EUR - AUS (A) | EUR - AUS (A) | NAm - novel (H10) | NAm - AUS | EUR - AUS (N3) | EUR - AUS (A) | A-EUR - novel |
| H10N3 | A/wild waterfowl/Western Australia/AS-22-0673-0006/2022(H10N3) | 16/2/2022 | na<sup>2</sup> | na<sup>2</sup> | na<sup>2</sup> | NAm - novel (H10) | NAm - AUS | EUR - AUS (N3) | EUR - AUS (C) | A-EUR - novel |
| H10N7 | A/chicken/New South Wales/M21-10080-0006/2021(H10N7) | 2/7/2021 | EUR - novel | EUR - novel | EUR - AUS (A) | NAm - novel (H10) | NAm - AUS | EUR - AUS (N7) | EUR - novel | A-EUR - novel |
| H10N7 | A/tawny frogmouth/Western Australia/21-1823/2021(H10N7) | 28/6/2021 | EUR - novel | EUR - novel | EUR - AUS (A) | NAm - novel (H10) | NAm - AUS | EUR - novel (N7) | EUR - AUS (A) | A-EUR - novel |
| H10N7 | A/Grey Teal/South Australia/22-64233564-17/2022(H10N7) | 21/5/2022 | EUR - AUS (A) | EUR - AUS (A) | EUR - AUS (A) | NAm - novel (H10) | NAm - AUS | EUR - AUS (N7) | EUR - AUS (A) | A-EUR - AUS (A) |
| H10N7 | A/Grey teal/Victoria/15848/2022(H10N7) | 21/6/2022 | EUR - AUS (A) | EUR - novel | EUR - AUS (A) | NAm - novel (H10) | NAm - AUS | EUR - AUS (N7) | EUR - novel | B-EUR-AUS |
| H10N7 | A/Grey teal/Victoria/15974/2022(H10N7) | 21/6/2022 | EUR - AUS (A) | EUR - novel | EUR - AUS (A) | NAm - novel (H10) | NAm - AUS | EUR - AUS (N7) | EUR - novel | B-EUR-AUS |
| H10N7 | A/Chestnut teal/Victoria/16047/2022(H10N7) | 28/6/2022 | EUR - AUS (A) | EUR - novel | EUR - AUS (A) | NAm - novel (H10) | NAm - AUS | EUR - AUS (N7) | EUR - novel | B-EUR-AUS |
| H10N7 | A/Chestnut teal/Victoria/16099/2022(H10N7) | 28/6/2022 | EUR - AUS (A) | EUR - novel | EUR - AUS (A) | NAm - novel (H10) | NAm - AUS | EUR - AUS (N7) | EUR - novel | B-EUR-AUS |
| H10N7 | A/Chestnut teal/Victoria/16148/2022(H10N7) | 5/7/2022 | EUR - novel | EUR - novel | EUR - AUS (A) | NAm - novel (H10) | NAm - AUS | EUR - AUS (N7) | EUR - novel | B-EUR-AUS |
| H10N7 | A/Grey teal/Victoria/16156/2022(H10N7) | 5/7/2022 | EUR - AUS (A) | EUR - novel | EUR - AUS (A) | NAm - novel (H10) | NAm - AUS | EUR - AUS (N7) | EUR - novel | B-EUR-AUS |
| H10N7 | A/Grey teal/Victoria/16164/2022(H10N7) | 5/7/2022 | EUR - AUS (A) | EUR - novel | EUR - AUS (A) | NAm - novel (H10) | NAm - AUS | EUR - AUS (N7) | EUR - novel | B-EUR-AUS |
| H10N7 | A/Grey teal/Victoria/16182/2022(H10N7) | 5/7/2022 | EUR - AUS (A) | EUR - novel | EUR - AUS (A) | NAm - novel (H10) | NAm - AUS | EUR - AUS (N7) | EUR - novel | B-EUR-AUS |
| H10N7 | A/Grey teal/Victoria/16190/2022(H10N7) | 5/7/2022 | EUR - AUS (A) | EUR - novel | EUR - AUS (A) | NAm - novel (H10) | NAm - AUS | EUR - AUS (N7) | EUR - novel | B-EUR-AUS |
| H10N7 | A/Grey teal/Victoria/16195/2022(H10N7) | 5/7/2022 | EUR - AUS (A) | EUR - novel | EUR - AUS (A) | NAm - novel (H10) | NAm - AUS | EUR - AUS (N7) | EUR - novel | B-EUR-AUS |

<sup>1</sup>Lineage information is as follows:
EUR-AUS: Established Australian clade in the Eurasian lineage
EUR – novel: Novel incursion from the Eurasian lineage
NAm – AUS: Established Australian clade in the North American lineage
NAm – novel: Novel incursion from the North American lineage
In cases where viruses fell into different North Australian lineages, we had appended an arbitrary letter in brackets (e.g. A, B, C) indicates that while viruses fell into Australian lineages, they fell into different lineages.
In the case of NS, whether the virus fell into the A or B allele is indicated prior to the lineage name. E.g. B-EUR-Aus.
Phylogenetic trees demonstrating sequence placement are presented in Figure 1-2, Figure S1-S3.
<sup>2</sup> PB2, PB1 and PA sequences were not recovered from A/wild waterfowl/Western Australia/AS-22-0673-0006/2022(H10N3) nor A/wild waterbird/South Australia/22-68204541-55/2022(H4N6).

The five H4N8 viruses isolated from Red-necked Stints (*Calidris ruficollis*) (n=4) in 2020 and one from a Sharp-tailed Sandpiper (*Calidris acuminata*) (n=1) in 2021 comprised segments from both recent introduction events (*i.e.* were novel incursions) and established lineages circulating in Australia. As all five viruses from Red-necked Stints were recovered from birds captured at the same sampling event it is unsurprising that the viruses had the same genome constellation (Table 1, Figure 1, Figure 1, S4). Notably, the virus recovered from the Sharp-tailed Sandpiper, captured approximately a year later, had all 8 segments fall into the same clade as those viruses from the Red-necked Stints, and the sequences for all segments were consistently _≥_99.9% similar to each other. Both the HA and NA segments comprised a novel incursion into Australia from Eurasia, the HA and NA diverging from the most closely related LPAIV sequences in databases with a mean dates of August 2018 (95% highest posterior density [HPD] Oct 2017-April 2019) and September 2018 (95% HPD Sept 2017-June 2019), respectively (Figure 1, Table S2). There was a time lag of approximately one year between the mean date of divergence from these reference sequences and the mean most recent common ancestor (MRCA) of the clade comprising the six shorebird sequences recovered in Australia. In contrast, the mean MRCA for the six virus glycoprotein gene sequences, particularly the N8 (April 2020, 95% HDP Nov 2018-Oct 2020) were very close to the sample collection date (December 2020), suggesting likely proliferation though the population following a single introduction into Australia (Figure 1, Table S2). For both the HA and NA segments, the most closely related LPAIV publicly available sequence sequences were viruses sampled from South Korean wild birds, although a general lack of publicly available sequence data prevented any firm conclusions being drawn with regards to the geographical source of the viral incursion from Asia.

The evolutionary histories of the internal segments of these six H4N8 viruses are more complex, but generally fall into clades dominated by viruses detected in Red-necked Stints (Figure S4). These clades are unusual in that they comprise exportation events from Australia back to Asia, which has not been seen with other clades (Wille et al. 2022). In four segments (PB2, PA, NP, M), the clades were first detected in Australian waterfowl, prior to entering shorebird populations. In the remaining segments, clades were detected in Asian waterfowl, prior to detection in shorebirds in Australia (Figure S4). The PB2, PA, NP, and NS clades have been circulating in shorebirds in Australia since ∼2012, whereas the PB1 and M clades have only been detected in Australian shorebirds more recently (since ∼2016). Exportation events from Australia to Eurasia include detections in Red-necked Stints in Japan (*e.g.* A/*Calidris ruficollis*/Hokkaido/12EY0172/2012(H4N7)), and more recently, A/wild bird/Fujian/24/2017(H2N6) (Figure S4). Based on BLASTn (https://blast.ncbi.nlm.nih.gov/Blast.cgi) analysis of the NA segment of A/*Calidris ruficollis*/Hokkaido/12EY0172/2012(H4N7) (GenBank Accession LC467226), this N7 is most similar to three viruses detected in Red-necked Stints in Australia (GenBank accessions: OL370713, OL370689 OL370697), suggesting that this NA segment was also exported from Australia. Overall, based on the genome constellations of the H4N8 viruses described here (Figure 1, S1, S4, Table 1), it is likely that reassortment occurred withing Australian shorebirds following the incursion of novel H4N8 viruses.

Between 2004-2017, the now waterfowl-associated H4 Australian lineage was detected in *Calidris* shorebirds in Australia, in addition to both *Calidris* shorebirds and gulls in Asia, including A/*Calidris ruficollis*/Hokkaido/12EY0172/2012(H4N7), demonstrating circulation in shorebirds with associated export events (Figure S1). However, unlike the internal segments, a cross-host order transmission event occurred followed by widespread circulation in waterfowl with no further detection of this H4 clade in shorebirds (however this may be affected by undersampling of Australian shorebirds).

### Recent incursion and widespread transmission of H10 viruses in Australian birds

The 16 H10 viruses sequenced in this study, comprising H10N2 (n=1), H10N3 (n=2), and H10N7(n=13), were collected from across multiple states in Australia, including Western Australia, South Australia, New South Wales and Victoria from 2020 to 2022. The viruses were recovered from both national active surveillance of wild birds as well as from passive or general surveillance (Table S1). Seven different genome constellations were recovered, with novel incursion events recorded for six segments. Only the PA and NP segments were exclusively from enduring Australian lineages (Figure 2, Figure S2-S3, Table 1).

**Figure 2.**
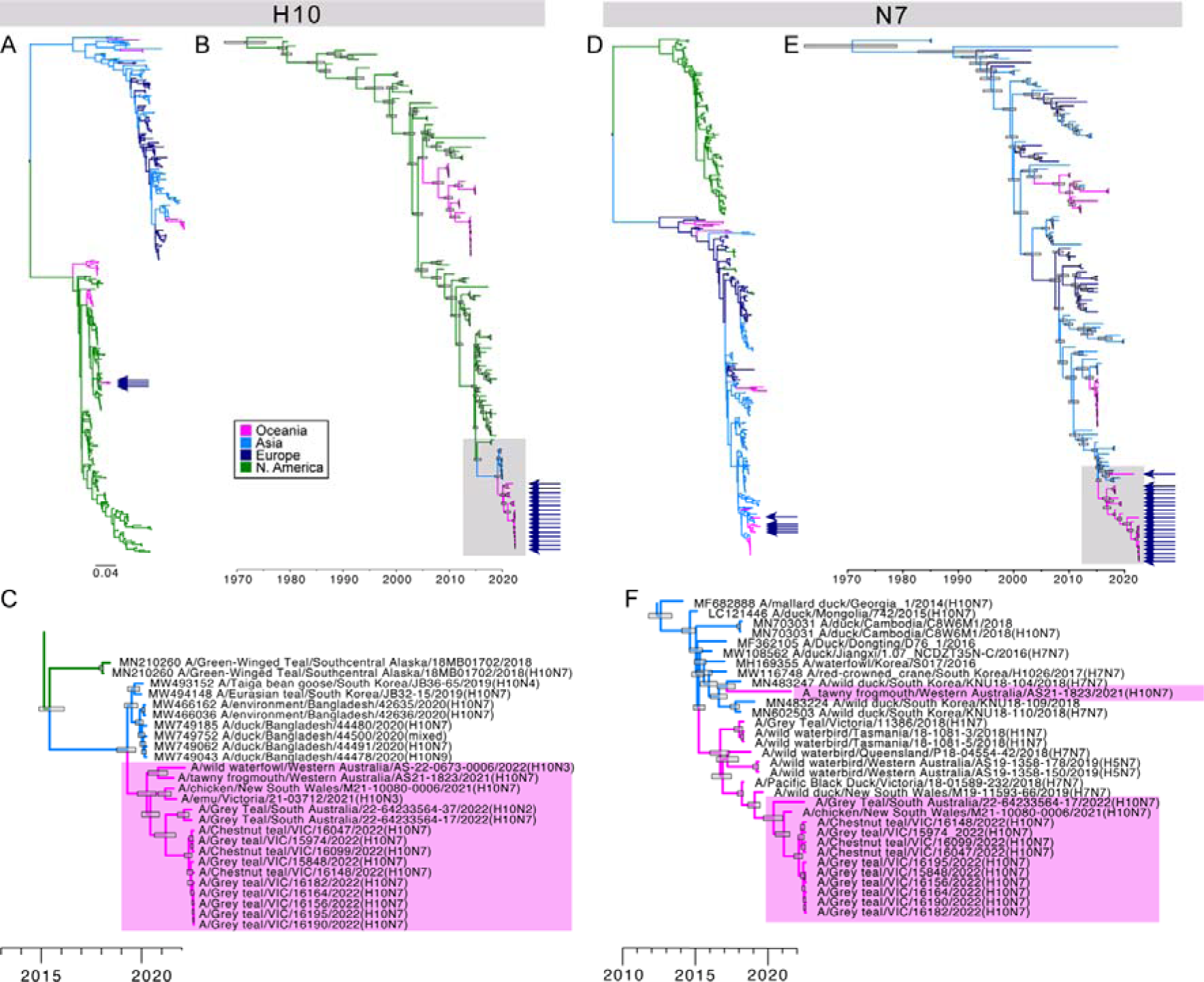
Phylogenetic trees of all (A,B,C) H10 sequences regardless of NA, and (D,E,F) N7 sequences. A phylogenetic tree for N2 and N3 is provided in Figure S2. (A, D) Maximum likelihood trees comprising all sequences collated for this study. Trees were rooted geographically, and the scale bar corresponds to number substitutions per site. (B, E) Time structured phylogenetic trees. The trees comprise select sequences from relevant lineages (here Eurasian lineage with all Australian sequences). (C, F), expansion of clade containing Australian sequences of interest, highlighted in a grey box in (B, E). Sequences of interest are indicated by arrows in (A, B, D, E) and are highlighted in a box in (C, F). Node bars correspond to the 95% highest posterior density (HPD) of node height.

All 16 HA sequences fell into the same H10 lineage (mean MRCA March 2020, 95% HPD 19 Sept 2019-20 Aug 2020). While these viruses fell into the broader North American clade of viruses, a viral incursion event in Asia occurred prior to their introduction in Australia, with detections in Bangladesh and South Korea in 2020 (Figure 2). Three different NA subtypes were detected, N2, N3 and N7 (Figure 2, S2, Table 1). Both N3 and the majority of the N7 sequences (12/13) fell into established Australian N3 and N7 lineages (Figure 2, S2, Table 1). A single N7 sequence (A/tawny frogmouth/Western Australia/21-1823/2021(H10N7)) and the N2 sequence (A/Grey Teal/South Australia/22-64233564-37/2022(H10N2)) represented novel incursions (Table 1, Figure S2, Figure 2), with the date of divergence of reference sequences January 2017 (95% HPD May 2016 - 18 Aug 2017) and December 2017 (95% HPD Jan 2017 - Oct 2018), respectively. In both cases there was a time lag of approximately 4 years between the date of divergence from the most closely related sequences in GenBank and detection in Australia, suggesting potential cryptic circulation for several years prior to detection (Table S2).

Focussing on the segments likely comprising novel incursions into Australia (PB2, PB1, HA, NA, M, NS) the mean MRCA estimates range from March – June 2020 (Figure S3, Table S2), and the dates of divergence from the closest sequences in reference databases range from 2019-2020. Among the H10 viruses, a number of different genome constellations were present, with a variable mix of novel and established Australian lineages, despite all sharing the same HA clade. This indicates different reassortment histories within Australia. For example, despite being detected on opposite sides of Australia and in different hosts and captivity settings, A/chicken/New South Wales/M21-10080-0006/2021(H10N7) and A/tawny frogmouth/Western Australia/21-1823/2021(H10N7) share a genome constellation across 6/8 segments, with differences in the NA and M segments (Figure 2, Figure S3, Table 1). Furthermore, while nine of the ten H10N7 viruses collected from *Anas* ducks at the same site in Victoria across 3 weeks in June and July 2022 share the same genome constellation, one virus, A/Chestnut teal/Victoria/16148/2022(H10N7) possessed a different novel Eurasian lineage PB2 segment demonstrating the potential for different reassortants to circulate in the same locations within the same host populations (Figure 2, Figure S3, Table 1).

## Discussion

We provide compelling evidence of shorebirds forming an important vector for inter-continental spread of avian influenza viruses between Asia and Australia and demonstrate key differences in the outcomes of viral introduction events resulting in contrasting opportunities for introduction, establishment, distribution and evolution of avian influenza viruses on the Australian continent.

The data generated here provide evidence for the recent incursion of two LPAIV subtypes into Australia, with contrasting patterns of establishment and spread following their introduction. Australia is a sink for viral diversity, such that avian influenza lineages circulate on the continent, in isolation, for many decades. However, viral incursions do occur as shown by the recent (2005-2015) MRCA of most HA lineages found in Australia, and that these lineages were unrelated to historic sequences (from the 1980’s) (Wille et al. 2022). Incursion events of H10 into Australia have been previously described, with epidemiology consistent with our findings. Specifically, an H10 virus from the North American lineage entered Australia in 2007/08, was detected in wild birds in Victoria and Tasmania, and was subsequently detected in chickens in Queensland and New South Wales in 2010 and 2012, respectively (Vijaykrishna et al. 2013, Hoye et al. 2021). Here, we describe a new introduction of North American lineage H10, with initial detections occurring within the space of two months (June-August 2021), in backyard domestic chicken, farmed emu and captive Tawny Frogmouth in a wildlife zoo from both western and eastern states of Australia. In addition to an expanded avian host range, H10 has caused outbreaks in numerous mammalian species including seals (Bodewes et al. 2015, Krog et al. 2015, Bodewes et al. 2016), mink (Berg et al. 1990), and humans (Arzey et al. 2012). Indeed, the H10 lineage described in 2012 in New South Wales (Vijaykrishna et al. 2013, Hoye et al. 2021) was also detected in poultry abattoir workers (Arzey et al. 2012). The human health risk posed by the current introduced avian H10 lineage in Australia is unknown but is expected to be low.

While it is likely that this novel H10 lineage is now established in Australia due to widespread detections, a viral introduction event does not necessarily result in the establishment of the lineage or uptake of all parts of its genome. Indeed, in the case of Australia where viruses are likely introduced by shorebirds, there are three possible outcomes: (1) viral introduction followed by extinction prior to detection or re-detection, (2) maintenance in shorebird hosts without transmission into other bird species, notably waterfowl, (3) transmission into the waterfowl population, resulting in either co-circulation or competitive exclusion of previously circulating viruses. There have been a number of instances of novel lineages detected in shorebirds, but with no evidence of their establishment. For example, Hurt *et al* (Hurt et al. 2006) described H11 viruses in Sharp-tailed Sandpipers detected in 2004, and a recent analysis of all Australian sequences demonstrated no other viruses have been detected in this lineage since (Wille et al. 2022).

In the invasive-species literature there is ample appreciation that the successful spread of an invader (or viral lineage) is preceded by the translocation, introduction and establishment of that invader, and that all four steps need to be completed effectively for the invasion to be successful (*e.g.* (Kolar et al. 2001)). In making risk assessments and in the development of mitigation strategies it is important to distinguish between these processes. Using LPAIV incursions into Australia by migratory shorebirds as an example, whether or not the translocation step is successful depends on the prevalence of the virus in the source population, the volume of migrants, and how it affects the host’s migration success (Risely et al. 2018). Importantly, successful introduction may depend on how the infection affects host fitness (*e.g.* illness leading to increased predation risk by dead-end hosts), the number, density and suitability of potential hosts at the location of introduction and the suitability of the new environment for the virus to survive outside the host. Whether the virus will establish at the new location depends on any temporal variation in these conditions, which may or may not result in an establishment bottleneck. For instance, densities of migratory shorebirds in Australia are very low during the Australian winter and high host-specificity of the virus will thus reduce its chance of establishment; with host-switching being a prerequisite for lineage establishment in Australia. Finally, successful invasion or spread of the virus to other parts of the continent also depends on how successful the virus is in surviving with the typical biotic and abiotic conditions in Australia, which may vary (dramatically) over time (Dalziel et al. 2016).

Evidence for the long-term maintenance of LPAIV exclusively in shorebirds without spread into waterfowl is provided by the internal segments of the H4N8 viruses described here. Not only have these internal segments been circulating in shorebirds for a number of years, but there is also evidence of exportation events with detection in *Calidris* shorebirds in Asia. A recent study demonstrated that viral exchange between shorebirds and ducks was uncommon relative to the rate of exchange within each host order (Hicks et al. 2022). This is reflected in phylogenetic studies of shorebird specific clades, such as those discrete clades of H11 and H12 viruses circulating in North American shorebirds (Wille et al. 2018). However, cross-order virus transmission does occur, as shown by the long-circulating H4 lineage detected in ducks. Historically this lineage has circulated in waterfowl (1980-2003, 2014-present) and shorebirds (2004-2017), resulting in a complex pattern of interspecies transmission and the propensity for viruses dispersed intercontinentally in shorebirds to enter waterfowl populations. Furthermore, viruses found in Ruddy Turnstones (*Arenaria interpres*) in Australia had genome constellations with complex origins – both geographically, but also in host origin – as a result of reassortment (Hoye et al. 2021, Wille et al. 2022). It is, however, important to take into consideration the low number of viral sequences generated each year, such that we may not have the power to detect cross-species transmission if they do not further transmit through populations.

Finally, there are two potential outcomes once viruses have emerged in wild waterfowl populations: co-circulation or competitive exclusion of the previously circulating viruses, as shown by H6 viruses in North America (Bahl et al. 2009). We suggest that the patterns we observed in H10 are consistent with competitive exclusion. That is, neither the H10 lineage reported by (Vijaykrishna et al. 2013, Hoye et al. 2021) nor an H10 lineage of Eurasian origin reported in (Wille et al. 2022) have been detected in wild birds since 2015 and 2019, respectively. Therefore, these lineages have likely been replaced by the novel H10 lineage reported here; alternatively they persist in locations or populations which have not been sampled adequately. In addition to lineage replacement through competitive exclusion, novel lineages reassort with locally circulating viruses resulting in genome constellations comprising segments that have been circulating in Australian wild birds for decades as well as segments recently introduced to Australia, as described both in this study, and elsewhere (Hoye et al. 2021, Wille et al. 2022). The timing of the H10 incursion may provide further insight into why viral spread, both spatially and across host species, was so frequent following emergence in waterfowl populations. Studies of LPAIV in Australia have demonstrated that high rainfall is associated with increased viral prevalence in wild birds (Ferenczi et al. 2016, Wille et al. 2023), which in turn is associated with outbreaks in poultry (Ferenczi et al. 2021) due to the high spillover potential from wild birds. In 2020 there was a shift to La Niña conditions in Australia, resulting in high rainfall following a number of years of drought (Meteorology 2023), leading to a dramatic increase in juvenile waterfowl.

The understanding of viral incursion and consequent dispersal is of critical importance for both LPAIV and HPAIV. The global HPAIV H5N1 panzootic is affecting wildlife on all continents except Australia (Oceania) (Klaassen et al. 2023, Wille et al. 2024). A greater understanding of the rates and risk factors for viral incursion, establishment and spread across space, time and host community of LPAIV, such as presented here, may play a key role in informing risk assessment and responses strategies for potential HPAIV incursions. Critically, our study highlights that shorebirds should be monitored for viral incursion, which will reveal whether the virus has established in the local waterfowl population (Wille et al. 2024). Studies of viral movement within Australia (Wille et al. 2022), as well as the understanding of environmental factors on LPAIV virus ecology in waterfowl (Wille et al. 2023) and spillover risk to poultry (Ferenczi et al. 2021) are critical to response planning.

## Acknowledgements

Surveillance and sampling performed by jurisdictional partners through the National Avian Influenza Wild Bird (NAIWB) Surveillance Program administered by Wildlife Health Australia, was supported with funding from the Australian Government Department of Agriculture, Fisheries and Forestry (DAFF). MK was supported by Australia Research Council Discovery grant (DP190101861). MW was funded by an ARC Discovery Early Career Researcher Award (DE200100977) (2020-2023). The WHO Collaborating Centre for Reference and Research on Influenza is supported by the Australian Government Department of Health. The Australian Centre for Disease Preparedness is supported by the Australian Government Department of Agriculture, Fisheries and Forestry. We are grateful to all those that contributed to sample collection.

---

**Table S1.**
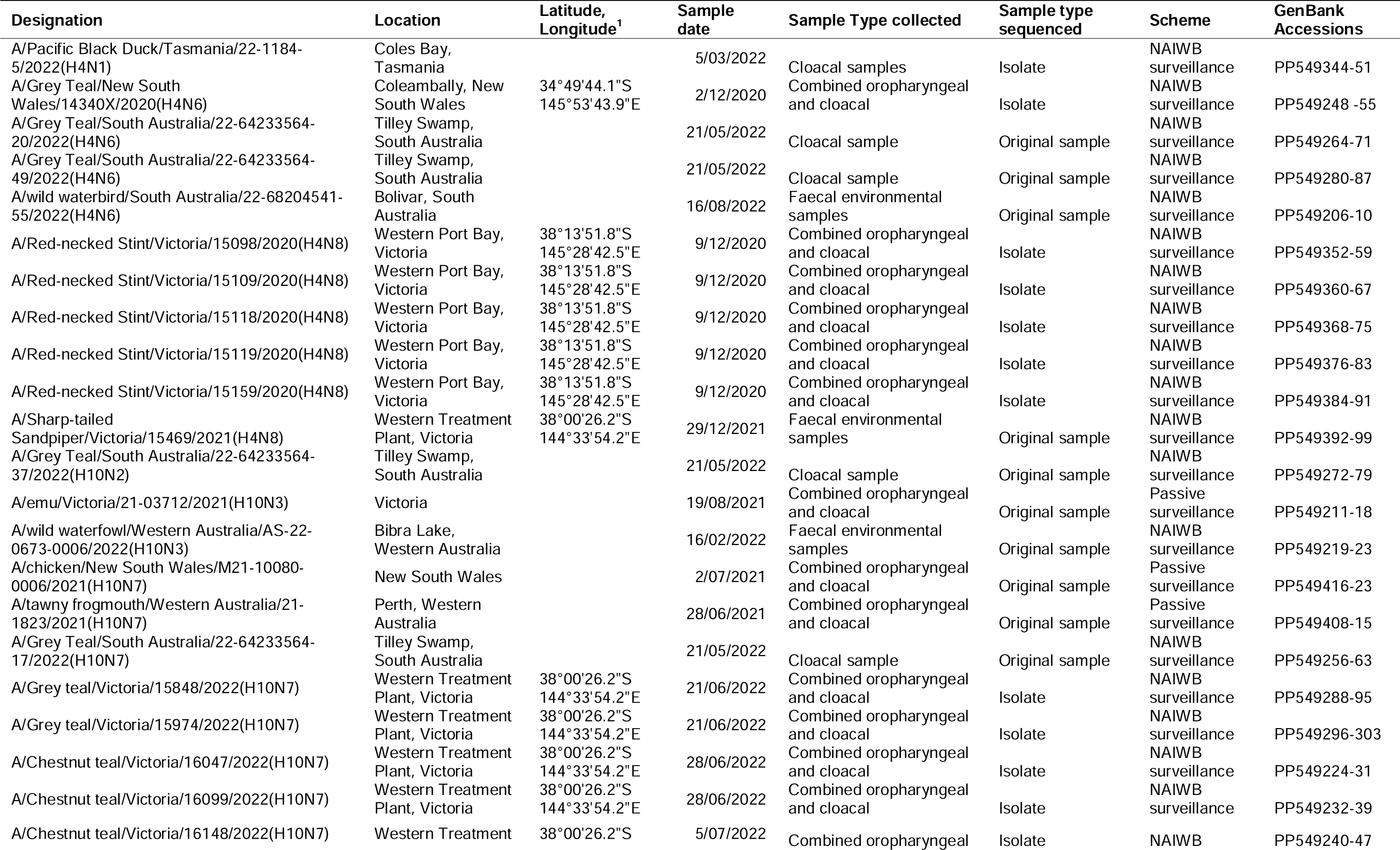
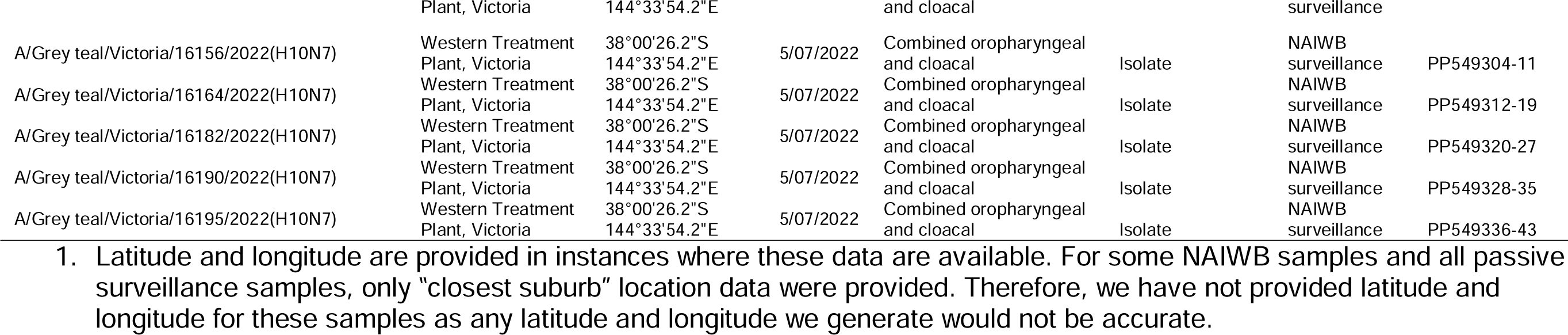
Metadata corresponding to H4 and H10 viruses included in this study.

**Table S2:**
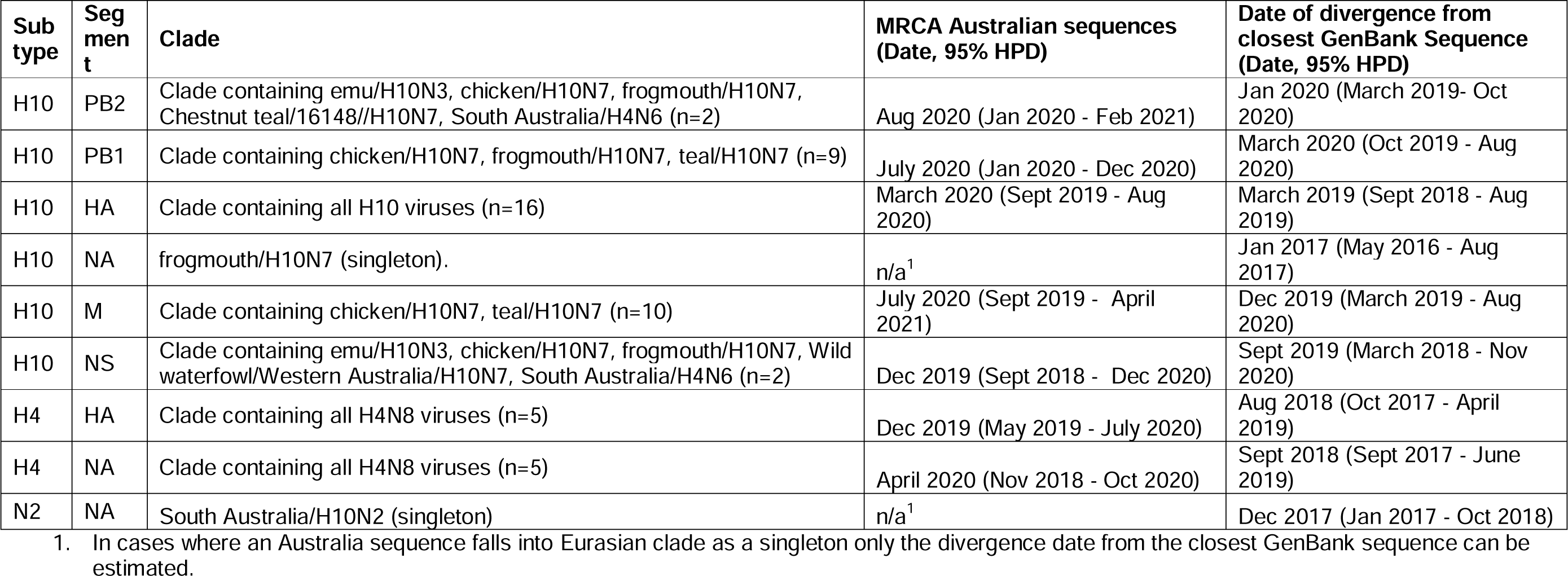
Estimated date of viral incursions into Australia for select segments and clades. Specific information pertaining to clade composition can be found in Figure 1-2, S2-S3, Table 1.

| Sub type | Segment | Clade | MRCA Australian sequences (Date, 95% HPD) | Date of divergence from closest GenBank Sequence (Date, 95% HPD) |
| --- | --- | --- | --- | --- |
| H10 | PB2 | Clade containing emu/H10N3, chicken/H10N7, frogmouth/H10N7, Chestnut teal/16148//H10N7, South Australia/H4N6 (n=2) | Aug 2020 (Jan 2020 - Feb 2021) | Jan 2020 (March 2019- Oct 2020) |
| H10 | PB1 | Clade containing chicken/H10N7, frogmouth/H10N7, teal/H10N7 (n=9) | July 2020 (Jan 2020 - Dec 2020) | March 2020 (Oct 2019 - Aug 2020) |
| H10 | HA | Clade containing all H10 viruses (n=16) | March 2020 (Sept 2019 - Aug 2020) | March 2019 (Sept 2018 - Aug 2019) |
| H10 | NA | frogmouth/H10N7 (singleton). | n/a<sup>1</sup> | Jan 2017 (May 2016 - Aug 2017) |
| H10 | M | Clade containing chicken/H10N7, teal/H10N7 (n=10) | July 2020 (Sept 2019 - April 2021) | Dec 2019 (March 2019 - Aug 2020) |
| H10 | NS | Clade containing emu/H10N3, chicken/H10N7, frogmouth/H10N7, Wild waterfowl/Western Australia/H10N7, South Australia/H4N6 (n=2) | Dec 2019 (Sept 2018 - Dec 2020) | Sept 2019 (March 2018 - Nov 2020) |
| H4 | HA | Clade containing all H4N8 viruses (n=5) | Dec 2019 (May 2019 - July 2020) | Aug 2018 (Oct 2017 - April 2019) |
| H4 | NA | Clade containing all H4N8 viruses (n=5) | April 2020 (Nov 2018 - Oct 2020) | Sept 2018 (Sept 2017 - June 2019) |
| N2 | NA | South Australia/H10N2 (singleton) | n/a<sup>1</sup> | Dec 2017 (Jan 2017 - Oct 2018) |

1. In cases where an Australia sequence falls into Eurasian clade as a singleton only the divergence date from the closest GenBank sequence can be estimated.

**Figure S1.**
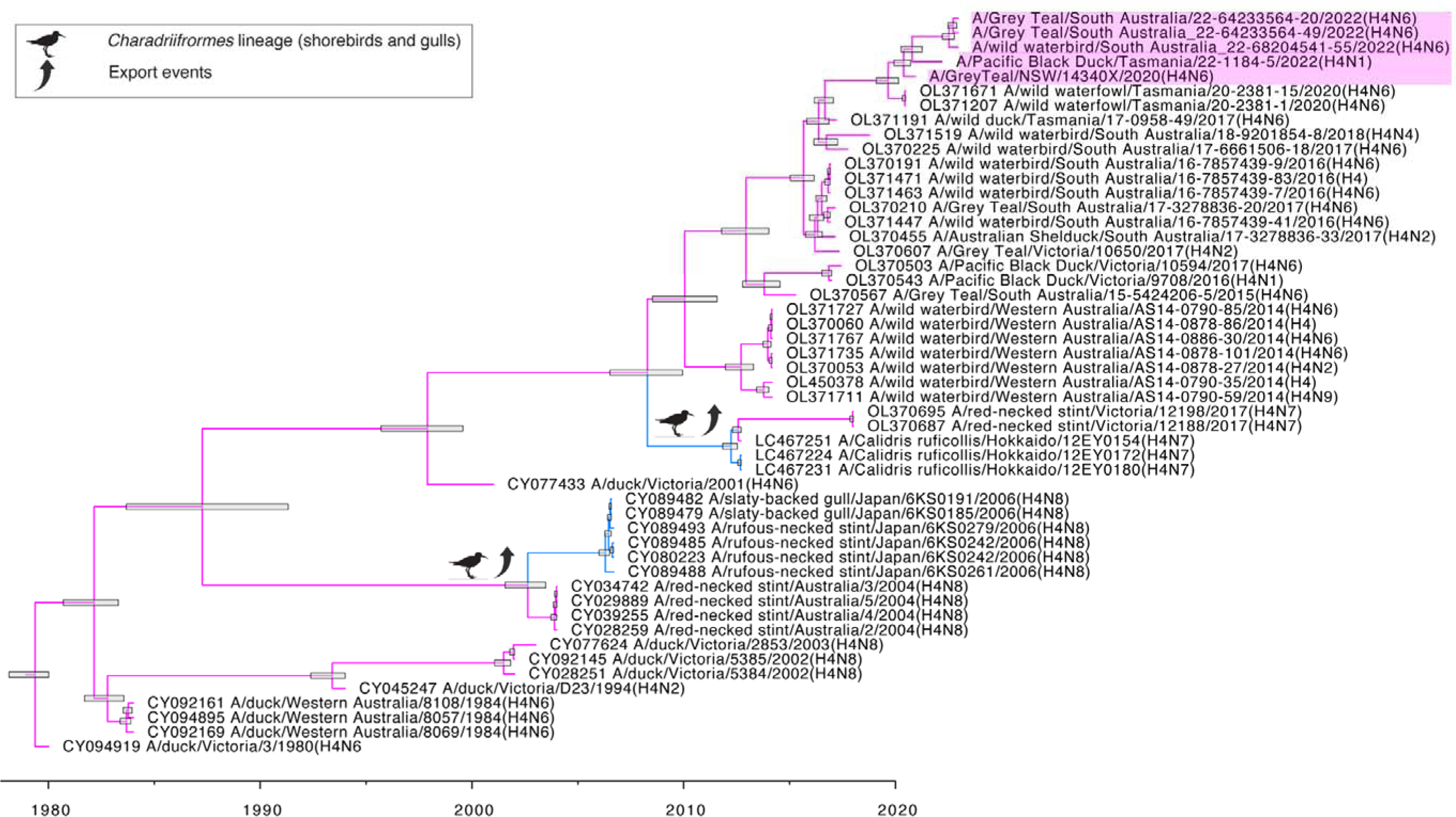
Expansion of H4 lineage from Figure 1 which has been circulating in Australia for four decades. Viruses isolated in this study are denoted by a pink box. Arrows indicate exportation events. Node bars correspond to the 95% highest posterior density (HPD) of node height. Branches are coloured by continent, consistent with Figure 1.

**Figure S2.**
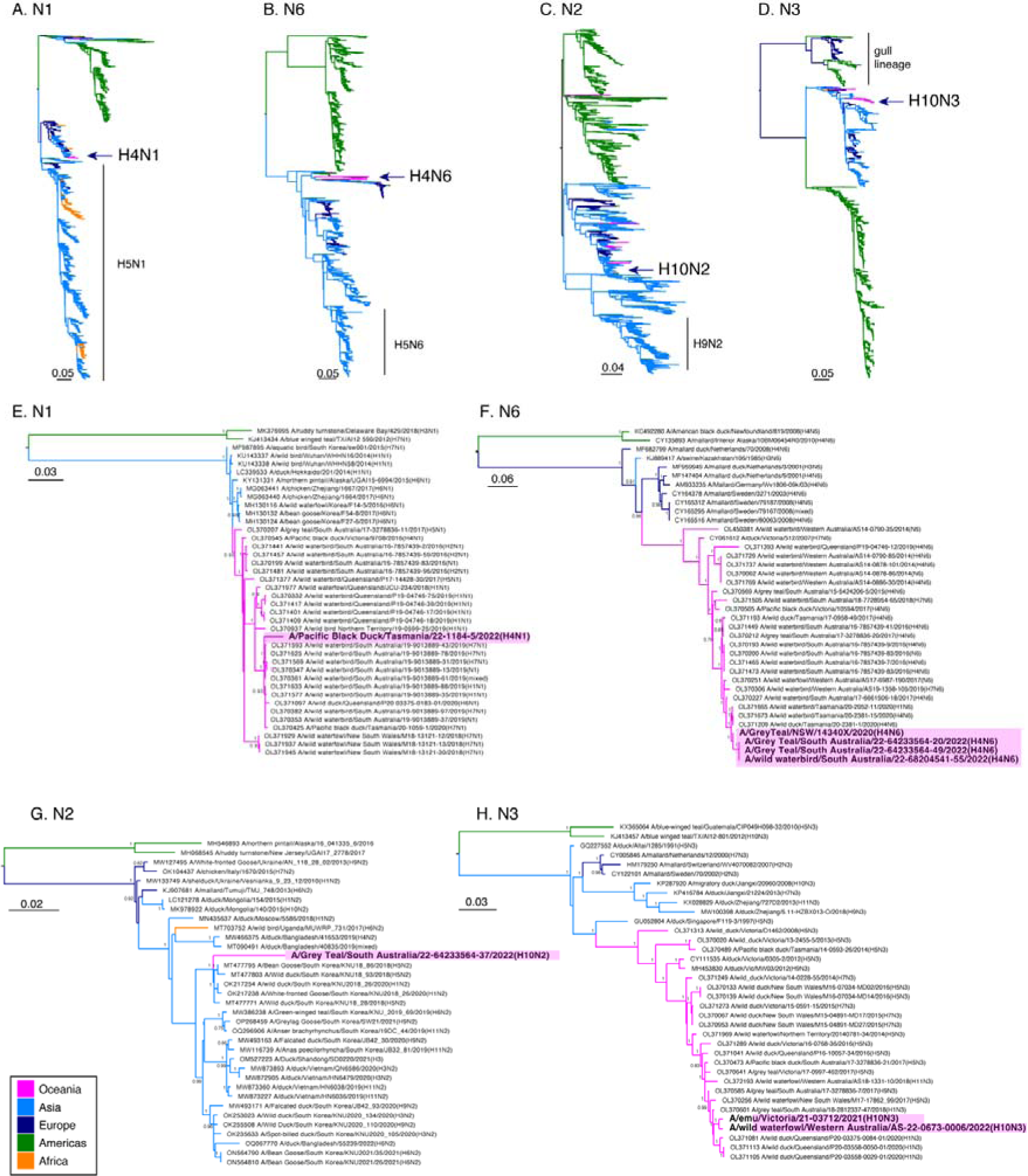
Phylogenetic trees of N1, N2, N6 and N3. (A, B, C, D) Maximum likelihood trees of all sequences collated for this study. The N1, N2 and N6 tree is rooted based on broad geographic clades, and N3 is rooted against the gull-specific clade. (E, F, G, H) Maximum likelihood trees of closest BLAST hits. Trees were rooted based on sequences from the North American clade Sequences generated in this study are indicated by an arrow in (A, B, C, D) and a filled box in (E, F, G, H). Node labels correspond posterior probabilities calculated by aBayes, and the scale bar indicates the number of substitutions per site.

**Figure S3.**
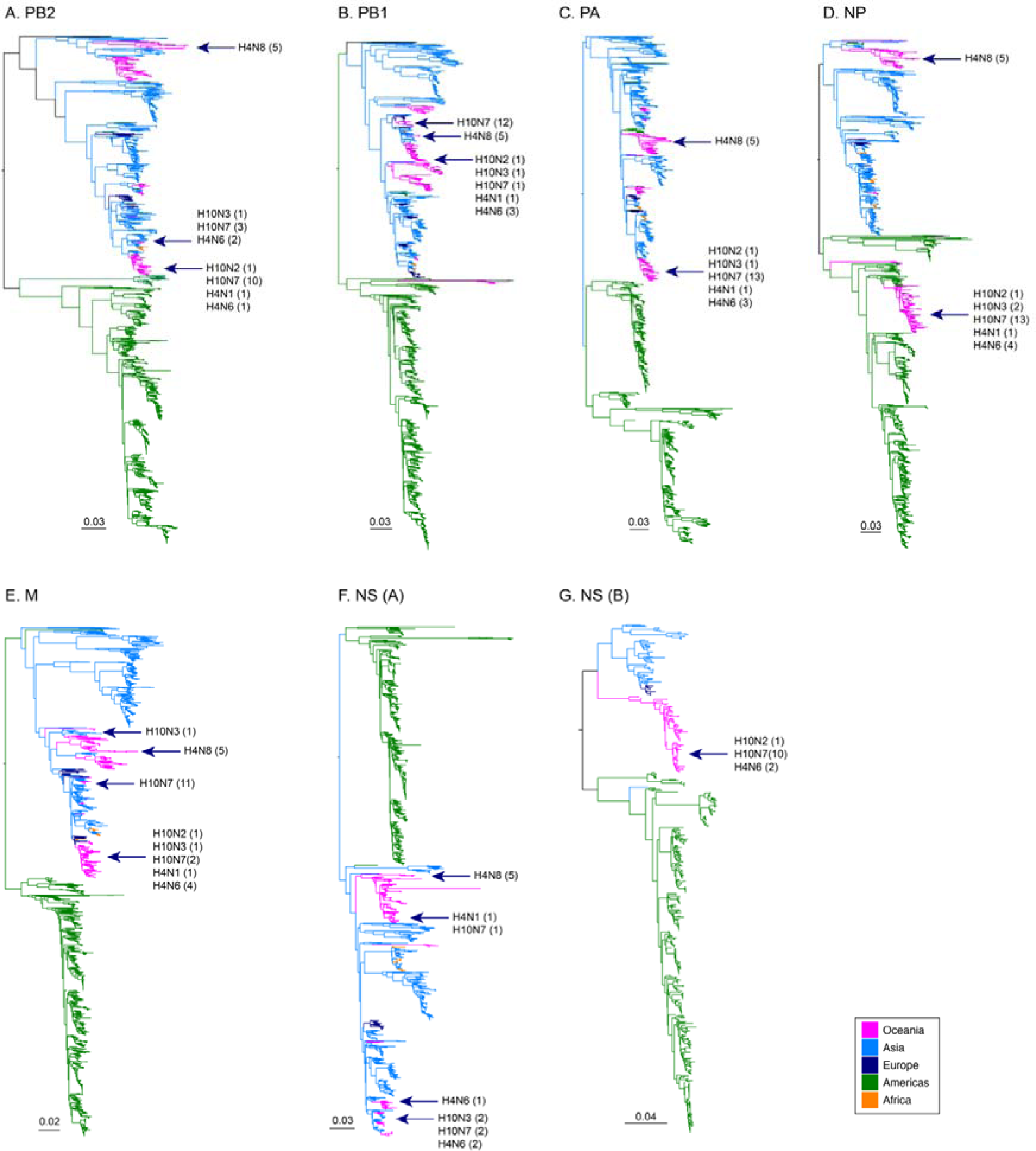
Phylogenetic trees for internal segments for all viruses sequenced in this study. Viruses are denoted by arrows and subtypes provided. Numbers in parentheses indicate the number of sequences. For the NS segment, the A and B allele were constructed separately. Trees were rooted based on the geographic division between American and Eurasian clades. The scale bar indicates the number of substitutions per site. Summary of clades presented in Table 1, and detailed phylogenies of H4N8 viruses presented in Figure S4.

**Figure S4.**
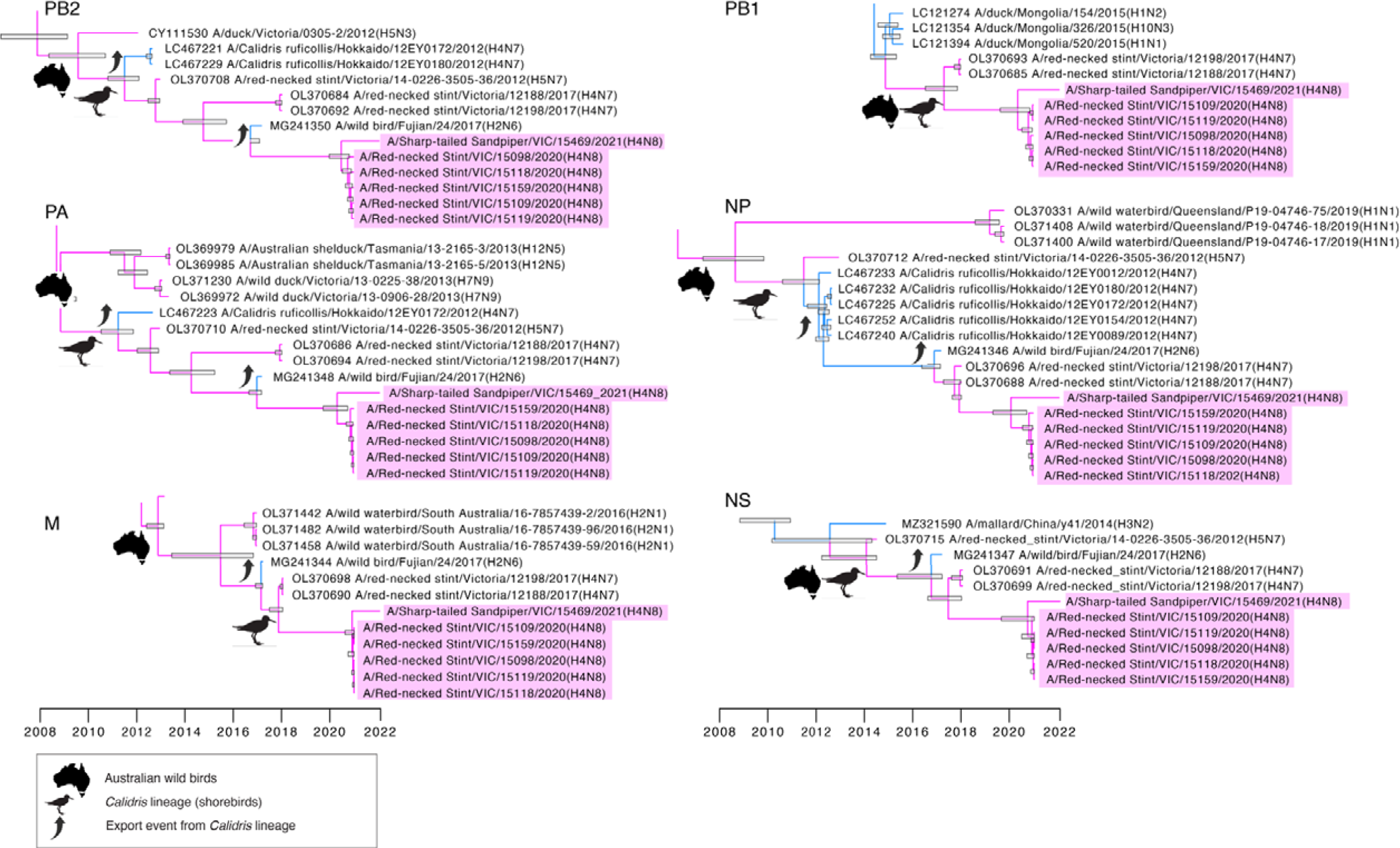
Time structured phylogenetic trees of internal segments of H4N8 viruses isolated from *Calidris* shorebirds. Viruses isolated in this study are denoted by pink boxes. Node bars correspond to the 95% HPD of node height. Branches are coloured by continent, consistent with Figure 1.

